# Characterisation of the carbapenem-resistant *Acinetobacter baumannii* clinical reference isolate BAL062 (CC2:KL58:OCL1): resistance properties and capsular polysaccharide structure

**DOI:** 10.1101/2024.05.09.593323

**Authors:** Alexander S. Shashkov, Nikolay P. Arbatsky, Sof’ya N. Senchenkova, Andrei S. Dmitrenok, Mikhail M. Shneider, Yuriy A. Knirel, Ruth M. Hall, Johanna J. Kenyon

**Affiliations:** N.D. Zelinsky Institute of Organic Chemistry, Russian Academy of Sciences, Moscow, Russia; M. M. Shemyakin & Y. A Ovchinnikov Institute of Bioorganic Chemistry, Russian Academy of Sciences, Moscow, Russia; School of Life and Environmental Science, The University of Sydney, Sydney, Australia; Centre for Immunology and Infection Control, School of Biomedical Sciences, Faculty of Health, Queensland University of Technology, Brisbane, Australia; School of Pharmacy and Medical Sciences, Health Group, Griffith University, Gold Coast, Australia

**Keywords:** *Acinetobacter baumannii*, BAL062, capsular polysaccharide, KL58, 8ePse, 5,7-diacetamido-3,5,7,9-tetradeoxynon-2-ulosonic acid

## Abstract

The carbapenem resistant *Acinetobacter baumannii* isolate BAL062 is a clinical reference isolate used in several recent experimental studies. It is from a ventilator associated pneumonia (VAP) patient in an intensive care unit at the Hospital for Tropical Diseases (HTD), Ho Chi Minh City, Vietnam in 2009. Here, BAL062 was found to belong to the B sub-lineage of global clone 2 (GC2) isolates in the previously reported outbreak (2008 and 2012) of carbapenem-resistant VAP *A. baumannii* at the HTD. While related sub-lineage B outbreak isolates were extensively antibiotic resistant and carry GC2-associated genomic resistance islands, AbGRI1, AbGRI2 and AbGRI3, BAL062 has lost AbGRI3 and three aminoglycoside resistance genes, *armA, aacA4* and *aphA1*, leading to amikacin and kanamycin susceptibility. The location of Tn*2008*VAR found in the chromosome of this sub-lineage was also corrected. Like many of the outbreak isolates, BAL062 carries the KL58 gene cluster at the capsular polysaccharide (CPS) synthesis locus and an annotation key is provided. As information about K type is important for development of novel CPS-targeting therapies, the BAL062 K58-type CPS structure was established using NMR spectroscopy. It is most closely related to K2 and K93, sharing similar configurations and linkages between K units and contains the rare higher monosaccharide, 5,7-diacetamido-3,5,7,9-tetradeoxy-d-*glycero*-l-*manno*-non-2-ulosonic acid (5,7-di-*N*-acetyl-8-epipseudaminic acid; 8ePse5Ac7Ac), the 8-epimer of Pse5Ac7Ac (5,7-di-*N*-acetylpseudaminic acid). Inspection of publicly available *A. baumannii* genomes revealed a wide distribution of the KL58 locus in geographically diverse isolates belonging to several sequence types that were recovered over two decades from clinical, animal, and environmental sources.

**IMPORTANCE:** Many published experimental studies aimed at developing a clearer understanding of the pathogenicity of carbapenem resistant *Acinetobacter baumannii* strains currently causing treatment failure due to extensive antibiotic resistance are undertaken using historic, laboratory adapted isolates. However, it is ideal if not imperative that recent clinical isolates are used in such studies. The clinical reference isolate characterized here belongs to the dominant *A. baumannii* GC2 clone causing extensively resistant infections, and has been used in various recent studies. Correlation of resistance profiles and resistance gene data is key to identifying genes available for gene knockout and complementation analyses, and we have mapped the antibiotic resistance genes to find candidates. Novel therapies, such as bacteriophage or monoclonal antibody therapies, currently under investigation as alternatives or adjuncts to antibiotic treatment to combat difficult-to-treat CRAb infections often exhibit specificity for specific structural epitopes of the capsular polysaccharide (CPS), the outer-most polysaccharide layer. Here, we have solved the structure of the CPS type found in BAL062 and other extensively resistant isolates. As consistent gene naming and annotation are important for locus identification and interpretation of experimental studies, we also have correlated automatic annotations to the standard gene names.

## INTRODUCTION

Carbapenem resistant *Acinetobacter baumannii* (CRAb) are a leading cause of antibiotic resistant nosocomial infections worldwide (1) and have limited treatment options remaining (2). Hence, alternate therapies are currently being sought. Although other clonal complexes (CC) such as CC1 (GC1), CC10, CC25 and CC79 are important, the clonal complex CC2 (also known as Global Clone 2, GC2) that is found on all inhabited continents, accounts for the majority of extensively resistant nosocomial *A. baumannii* isolates.

Owing to concerns about the use of early *A. baumannii* isolates such as ATCC17978 and ATCC19606 to study the pathogenesis of *A. baumannii*, particularly that they may be laboratory adapted and hence not strictly representative of current clinical isolates, a number of clinical isolates have begun to be used (3–5). The *A. baumannii* isolate BAL062 is a clinical carbapenem resistant GC2 isolate that has been utilised for this purpose. BAL062 had been recovered in 2009 from a patient with ventilator associated pneumonia (VAP) in an intensive care unit (ICU) at the Hospital for Tropical Diseases (HTD) in Ho Chi Minh City, Vietnam (6). It has since been used to develop a TraDIS library (7) and the complete genome sequence is available (NCBI GenBank accession number LT594095.1; (8)). BAL062 has also been used as a clinical reference isolate in several experimental studies. The BAL062 library has been used to identify genes that contribute to resistance to the last-resort antibiotic colistin (8, 9), and several clinically relevant biocides (7, 10). Additional studies on spermidine/spermine efflux (11), a comparison to other clinical and environmental isolates (12, 13), and demonstration of the utility of novel suicide vectors (14) have also used BAL062.

Previously, a series of carbapenem resistant isolates belonging to both GC2 and CC10 were reported to have caused an outbreak between 2008 and 2012 in the same ICU at the HTD in Ho Chi Minh City (15). Most of the carbapenem resistant isolates carried *oxa23*, the dominant and most widespread gene attributed to the spread of carbapenem resistance (16). However, *oxa23* is found in several distinct contexts (16, 17) and, if on the chromosome, their location can be characteristic for a specific lineage (18). Phylogenetic analysis revealed that the GC2 outbreak isolates could be separated into several distinct sub-lineages, designated A-E, and each sub-lineage had acquired *oxa23* independently. Lineage B carried a novel *oxa23*-containing transposon designated Tn*2008VAR* (Fig. 1A). Only lineage D carried KL2 at the chromosomal K locus (KL) for biosynthesis of the capsular polysaccharide (CPS) and KL2 was believed to be ancestral. Most lineage E carried KL49, and lineages A-C isolates carried KL58, with a single exception where KL32 had replaced KL58 (15). Recently, four HTD GC2 isolates from this outbreak were compared with ATCC17978 and shown to have increased virulence in mice with systemic dissemination and persistent colonisation of airways (19). This included two isolates with KL58 (BAL084 lineage B; BAL215 lineage C), one with KL2 (BAL276 lineage D) and one with KL49 (BAL191 lineage E),

**Figure 1.**
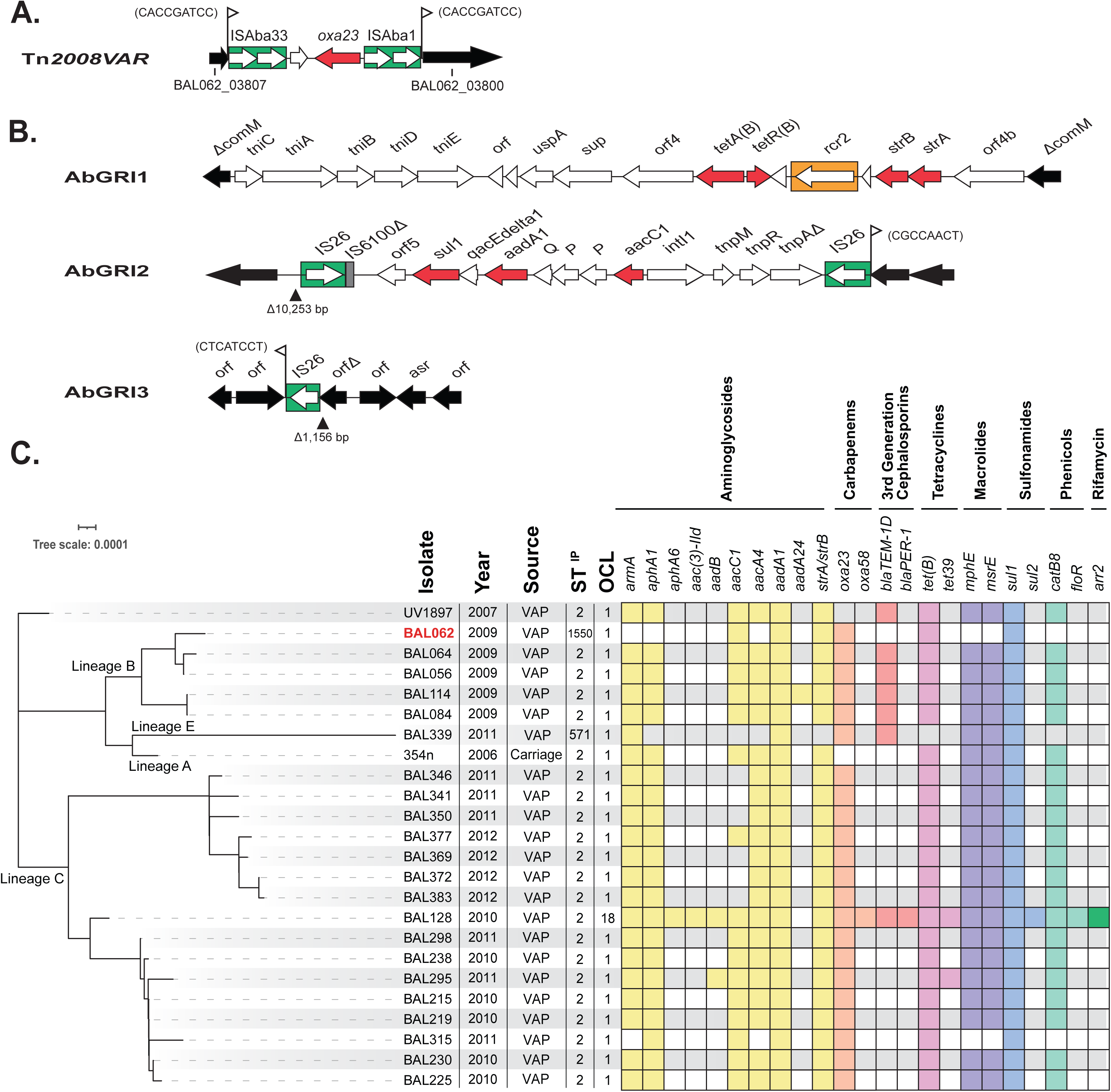
**(A)** Tn*2008VAR* in the BAL062 chromosome (base positions 3893121-3898596). The locus tags indicated on either side are the remnants of the interrupted acyl-CoA dehydrogenase gene in the chromosome. The sequence of the 9 bp target site duplication is shown next to the flags. **(B)** Genetic arrangement of AbGRIs in BAL062 chromosome: AbGRI1 (base positions 3779179-3801151); AbGRI2 (base positions 2675982-2686299); AbGRI3 (base positions 1400803-1408181). Green boxes indicate insertion sequences, red are resistance genes, orange box is CR2, and flanking chromosomal genes are black. **(C)** Core-SNP maximum likelihood phylogeny of GC2 genomes carrying KL58 from Vietnam HTB outbreak reported in Schultz et al. (SRA accession numbers listed in Table S1). BAL062 is shown in red. Year of collection, isolation source, ST^IP^ and OCL for each isolate are shown next to a presence/absence matrix of antibiotic resistance genes coloured by class. Lineages indicated in Schultz et al. are indicated.

Though the Vietnam GC2 outbreak isolates were resistant to a number of antibiotics, and the *oxa23* carbapenem resistance gene was present in different contexts, the remaining antibiotic resistance genes were not examined or reported previously (15). Most carbapenem resistant GC2 isolates carry chromosomal islands known as AbGRI1 and AbGRI2 that include genes generally conferring resistance to early antibiotics (20–23). A third, chromosomally-located resistance island, AbGRI3, that includes the *armA* gene is not found in early GC2 isolates, but is present in many isolates recovered after 2003 (24). The *armA* gene confers resistance to all clinically relevant aminoglycosides which are used as a last resort to treat carbapenem resistant infections (25).

Though not included in the previous study, BAL062 is clearly from the same outbreak granted its place and year of isolation. Here, we have placed BAL062 within one of the specific sub-lineages identified previously, and examined the resistance gene profiles of all the GC2 outbreak isolates. BAL062 was found to carry KL58, and we also report the structure of the K58 type CPS produced by BAL062.

## Results

### *A. baumannii* BAL062 is a multiply antibiotic resistant GC2 isolate

The complete genome sequence of BAL062 (NCBI GenBank accession numbers LT594095.1 (chromosome) and LT594096.1 (plasmid)) indicates that it belongs to sequence type (ST) 1550 in the *A. baumannii* Institut Pasteur (IP) multi-locus sequence typing (MLST) scheme, identifying it as a single locus variant (SLV) of ST2 that represents GC2. This was also noted recently (12).

Previously, BAL062 had been recorded as resistant to carbapenems (imipenem), penicillins and β-lactamase inhibitors (piperacillin/tazobactam, ampicillin), fluroquinolones (ofloxacin), third generation cephalosporins (ceftazidime, ceftriaxone, cefepime), aminoglycosides (gentamicin and amikacin), and sulfonamides and trimethoprim (co-trimoxazole) (9). Analysis of antibiotic resistance determinants revealed that resistance to carbapenems was due to the presence of an *oxa23* gene (locus tag BAL062_03803) within an unusual Tn*2008-*like transposon, previously designated Tn*2008VAR* (15). Tn*2008VAR* interrupted an acyl-CoA dehydrogenase gene in the chromosome generating a 9 bp target site duplication (Fig. 1A) and this location supersedes the location proposed originally. This transposon was previously found only in the B sub-lineage of the KL58 monophyletic clade (15). An appropriately oriented ISAba1 upstream of the *ampC* gene (locus tag BAL062_01109) accounts for resistance to third generation cephalosporins. Mutations in the quinolone-determining region of GyrA and ParC explain the fluoroquinolone resistance. However, a determinant for amikacin resistance was not found.

The genome also includes *strA-strB* for spectinomycin resistance and *tet(B)* for tetracycline resistance, which are both located in an AbGRI1-type island in the *comM* gene (Fig. 1B). This island is a Tn*6022*-derived transposon carrying a complete set of transposition genes (*tniC-tniA-tniB-tniD-tniE*). The *sul1* (sulfonamide resistance), *aadA1* (streptomycin and spectinomycin resistance), and *aacC1* (gentamicin resistance) genes are located in an IS*26*-bounded AbGRI2 type island (Fig. 1B). However, only an IS*26* remains of the IS*26*-bounded AbGRI3 island (Fig. 1B) suggesting that the AbGRI3 resistance genes had been lost during storage of the original isolate.

Therefore, the antibiotic susceptibility of BAL062 was re-evaluated using an extended panel of antibiotics. This showed that BAL062 was indeed susceptible to amikacin, as well as to tobramycin and kanamycin. It was also resistant to tetracycline and further resistant to meropenem and doripenem (carbapenems), ciprofloxacin and nalidixic acid, consistent with the resistance gene profile determined for this isolate.

### *A. baumannii* BAL062 is a member of GC2:KL58 sub-lineage B

The BAL062 genome was found to include the KL58 sequence at the CPS biosynthesis K locus (base positions 3946059 to 3973073) and OCL1 at the OC locus (base positions 587145 to 598627) that determines the outer-core (OC) structure of the lipooligosaccharide. During the HTD outbreak, the KL58 locus had been identified in 29 isolates belonging to either GC2 (n=23) or CC10 (n=6) ((15); Table S1). To assess the relationship of BAL062 to the GC2:KL58 HTD outbreak isolates, a core-SNP phylogeny was constructed (Fig. 1C). In this phylogeny, BAL062 was positioned within the B sub-lineage, which included four ST2 isolates, BAL056, BAL064, BAL084 and BAL114, that were recovered in the same year (2009) and had been reported to include the Tn*2008VAR* transposon (15).

The additional antibiotic resistance determinants detected were mapped against the tree and, while the other B isolates included many of the resistance genes found in BAL062, consistent with the presence of AbGRI1 and AbGRI2, they also carried *armA, aphA1* and *aacA4* aminoglycoside resistance genes, as well as *bla*_TEM-1D_*, mphE*-*msrE* and *catB8* genes that were absent from the BAL062 genome (Fig. 1C). This confirmed that the BAL062 isolate currently being used and used to determine the draft (9) and complete (8) genomes had lost the resistance genes expected to be present in AbGRI3 and some of those generally associated with AbGRI2.

### KL58 is related to KL2 and KL93

Annotations for the KL58 sequence are available in the BAL114 KL58 sequence record under GenBank accession number KT359617.1, and this sequence is 100% identical (100% coverage) to KL58 in the BAL062 genome (locus tags BAL062_03872-BAL062_03850). As consistent annotation is key to recognizing the function of genes identified in experimental studies, in Table 1 the standard annotations for *A. baumannii* K loci (26–28) that are used in most publications are compared to those generated using Prokka (29) that appear on the BAL062 genome (LT594095.1) and most automatically annotated genomes. Table 1 also includes the standard and automatic annotations for OCL1 (30, 31).

**Table 1.** Updated gene annotations for the KL58 and OCL1 loci in the BAL062 genome.

| Gene name | Locus tag | GenPept accession | Annotation in LT594095.1 | Function/Predicted Function |
| --- | --- | --- | --- | --- |
| **KL58 locus** |  |  |  |  |
| *wzc* | BAL062_03872 | SBS23916.1 | *ptk* | Protein tyrosine kinase |
| *wzb* | BAL062_03871 | SBS23915.1 | *ptp* | Low molecular weight protein tyrosine phosphatase |
| *wza* | BAL062_03870 | SBS23914.1 |  | Outer membrane protein |
| *gna* | BAL062_03869 | SBS23913.1 | *tuaD_2* | UDP-N-acetyl-galactosamine dehydrogenase |
| *psaA* | BAL062_03868 | SBS23912.1 | *capD* | UDP-N-acetylglucosamine 4,6-dehydratase/5-epimerase |
| *psaB* | BAL062_03867 | SBS23911.1 | *arnB* | C4-aminotransferase |
| *psaC* | BAL062_03866 | SBS23910.1 | *neuA* | Cytidylyltransferase |
| *psaD* | BAL062_03865 | SBS23909.1 |  | Nucleotidase |
| *psaE* | BAL062_03864 | SBS23908.1 |  | N-acetyltransferase |
| *psaF* | BAL062_03863 | SBS23907.1 | *spsE* | Condensase |
| *wzx* | BAL062_03862 | SBS23906.1 |  | Oligosaccharide-unit translocase |
| *gtr118* | BAL062_03861 | SBS23905.1 | *lst* | Glycosyltransferase |
| *wzy* | BAL062_03860 | SBS23904.1 |  | Oligosaccharide-unit polymerase |
| *gtr8* | BAL062_03859 | SBS23903.1 | *tagE* | Glycosyltransferase |
| *gtr9* | BAL062_03858 | SBS23902.1 | *lsgF* | Glycosyltransferase |
| *itrA2* | BAL062_03857 | SBS23901.1 | *wcaJ* | GalNAc-1P initiating transferase |
| *galU* | BAL062_03856 | SBS23900.1 | *galU* | UDP-glucose-1-phosphate uridylyltransferase |
| *ugd* | BAL062_03855 | SBS23899.1 | *tuaD_1* | UDP-glucose 6-dehydrogenase |
| *gpi* | BAL062_03854 | SBS23898.1 | *pgi* | glucose-6-phosphate isomerase |
| *gne1* | BAL062_03853 | SBS23897.1 | *galE_2* | UDP-glucose/UDP-N-acetyl-glucosamine 4-epimerase |
| *atr42* | BAL062_03852 | SBS23896.1 |  | Acetyltransferase |
| *atr43* | BAL062_03851 | SBS23895.1 |  | Acetyltransferase |
| *pgm* | BAL062_03850 | SBS23894.1 | *manB* | Phosphoglucomutase/phosphomannomutase |
| **OCL1 locus** |  |  |  |  |
| *gtrOC1* | BAL062_00583 | SBS20708.1 |  | Glycosyltransferase |
| *gtrOC2* | BAL062_00584 | SBS20709.1 |  | Glycosyltransferase |
| *pda1* | BAL062_00585 | SBS20710.1 | *icaB* | Polysaccharide deacetylase |
| *gtrOC3* | BAL062_00586 | SBS20711.1 | *lpsC* | Glycosyltransferase |
| *gtrOC4* | BAL062_00587 | SBS20712.1 |  | Glycosyltransferase |
| *orf1 (ghy)* | BAL062_00588 | SBS20713.1 |  | Unknown |
| *gtrOC5* | BAL062_00589 | SBS20714.1 |  | Glycosyltransferase |
| *gtrOC6* | BAL062_00590 | SBS20715.1 |  | Glycosyltransferase |
| *gtrOC7* | BAL062_00592 | SBS20717.1 | *sacB* | Glycosyltransferase |

KL58 (Fig. 2) has an arrangement typical of all other sequences found at the K locus in *A. baumannii* genomes to date (26–28), in that it includes a central region that determines the specific CPS type flanked by a module of *wza-wzb-wzc* genes for CPS export and *galU-pgm* genes for synthesis of common sugar precursors. In the previous study, it was reported that the KL58 sequence carried by GC2 HTD outbreak isolates in sublineages A-C (Fig. 1C) had likely arisen via a 24 kb sequence replacement involving part of the KL2 locus that was imported from a CC10 KL58 isolate (15). The portion shared by KL2 and KL58 (Fig. 2) includes a module of *psaABCDEF* genes for the synthesis of the monosaccharide 5,7-di-*N*-acetylpseudaminic acid (Pse5Ac7Ac), which is a constituent found in the oligosaccharide K-units that make up the K2 CPS (32, 33).

**Figure 2.**
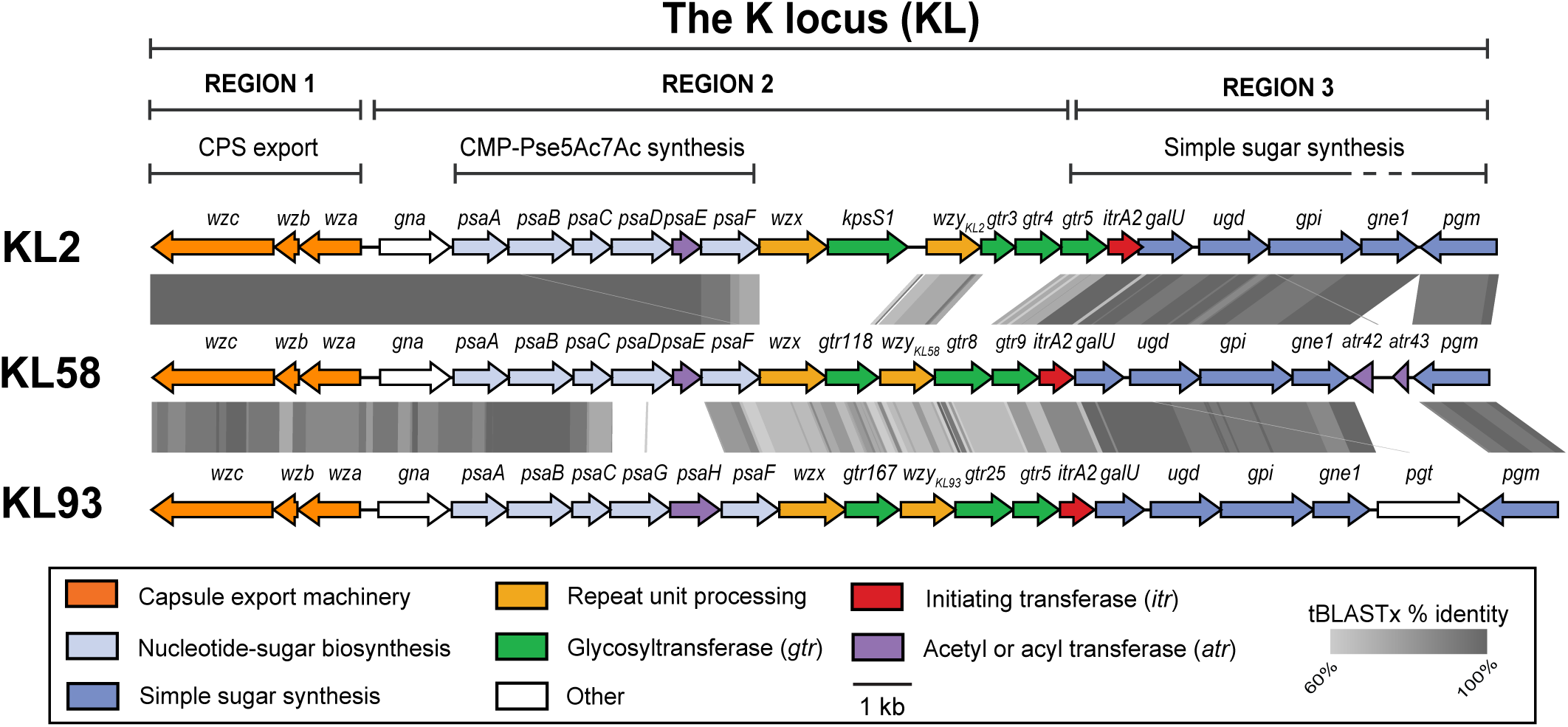
Comparison of KL58 in the BAL062 chromosome (base positions 3972158-3948095) with KL2 from *A. baumannii* A74 (GenBank accession number KJ459911) and KL93 from *A. baumannii* B11911 (GenBank accession number CP021345.1; bases 3338181-3368604). Genes coloured by function of gene product and grey shading is tBLASTx identity. Colour scheme and scale shown below.

The two loci differ in the region that includes predicted glycosyltransferase (*gtr*) genes and the Wzy polymerase gene for forming glycosidic linkages in the CPS, suggesting that the K2 and K58 structures are composed of similar monosaccharides that are linked together differently. This central portion in KL58 (*wzx-gtr9*) shares a level of sequence identity (>60% tBLASTx identity) with the *A. baumannii* KL93 sequence (Fig. 2), and as the K93 structure is related to K2 (34), the K58 structure is likely related to both CPS types. As the structure of the K58 type CPS is unknown, the structure of the CPS produced by BAL062 was determined.

### Monosaccharide composition of CPS recovered from BAL062

CPS was isolated from BAL062 cells and purified by Sephadex G-50 Superfine gel chromatography (see methods) for monosaccharide and structural analyses. Sugar analysis of the BAL062 CPS by GLC of the alditol acetates revealed the presence of glucose (Glc), galactose (Gal), galactosamine (GalN) and a higher order nonulosonate. The presence of signals for *N-*acetyl groups in the NMR spectra of the CPS (δ_С_ 23.0-23.8 (CH_3_) and 175.5–175.8 (CO), δ_H_ 2.00-2.10) indicated that all amino sugars are *N*-acetylated. Additional chemical analyses on the nonulosonate present revealed the sugar to be the 8-epimer of 5,7-*N*-acetylpseudaminic acid (Pse5Ac7Ac), known as 8ePse5Ac7Ac or 5,7-*N*-acetyl-3,5,7,9-tetradeoxynon-2-ulosonic acid. This sugar had only recently been discovered in the CPS of *A. baumannii* isolate RES-546 that carries the KL135 locus (35) and had not been described for any other isolate to date.

### Structural resolution of the CPS

To confirm the order of monosaccharides and overall topology of the BAL062 CPS, the complete structure was established by NMR spectroscopy (Fig. 3) using a set of shift-correlated two-dimensional NMR experiments (^1^H,^1^H COSY, ^1^H,^1^H TOCSY, ^1^H,^1^H ROESY, ^1^H,^13^C HSQC, and^1^H,^13^C HMBC). The spin-systems were revealed for the constituent monosaccharides, all being in the pyranose form. The chemical shifts of the monosaccharides are tabulated in Table 2, and the CPS structure is shown in Fig. 4.

**Figure 3.**
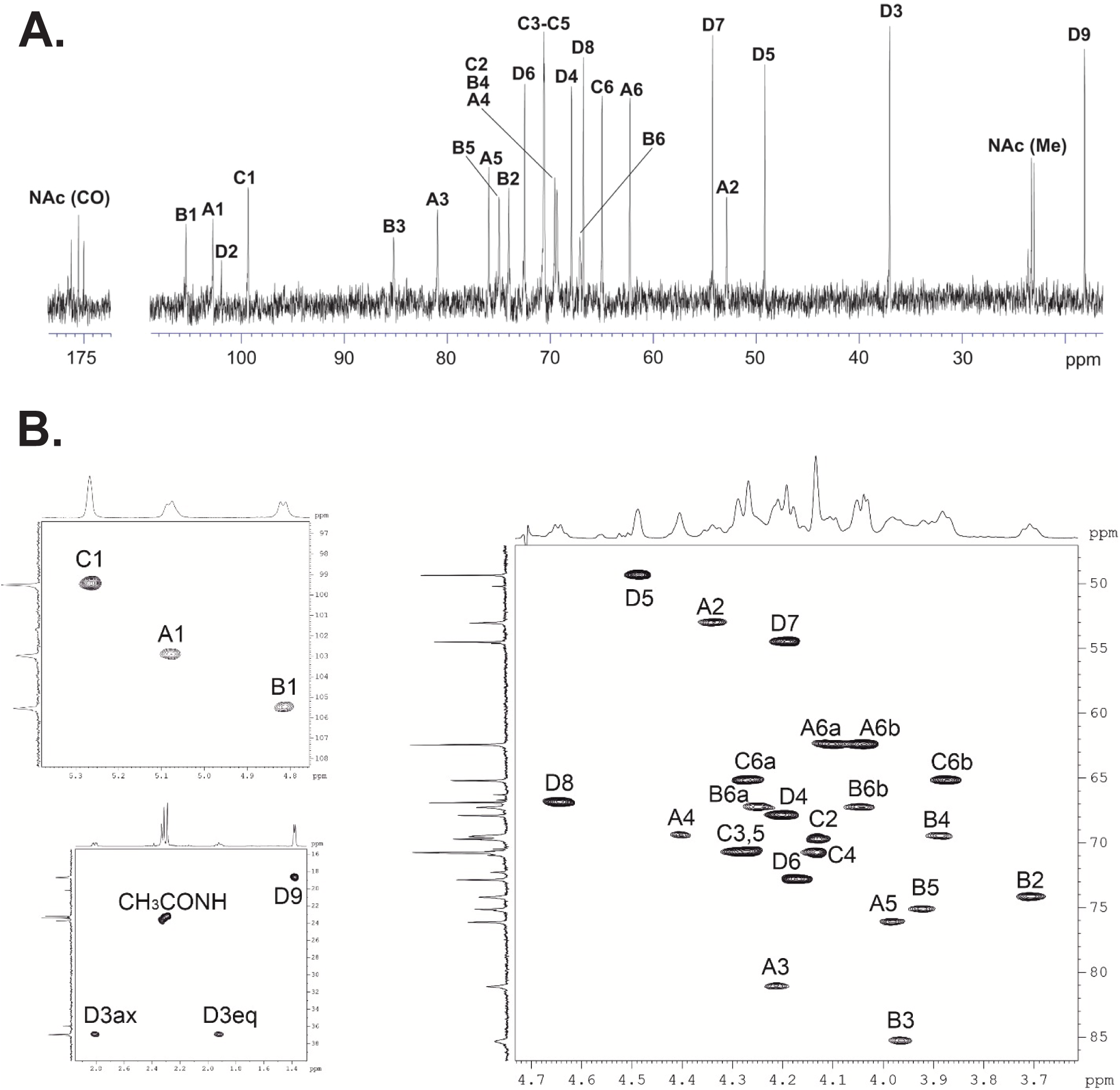
**(A)** ^13^C NMR spectra of the CPS of *A. baumannii* BAL062. **(B)** Parts of a two-dimensional ^1^H,^13^C HSQC spectrum of the CPS of *A. baumannii* BAL062. The corresponding parts of the one-dimensional ^1^H and ^13^C NMR spectra are displayed along the axes. For designations of the monosaccharide residues see Figure 4 and Table 2.

**Figure 4.**
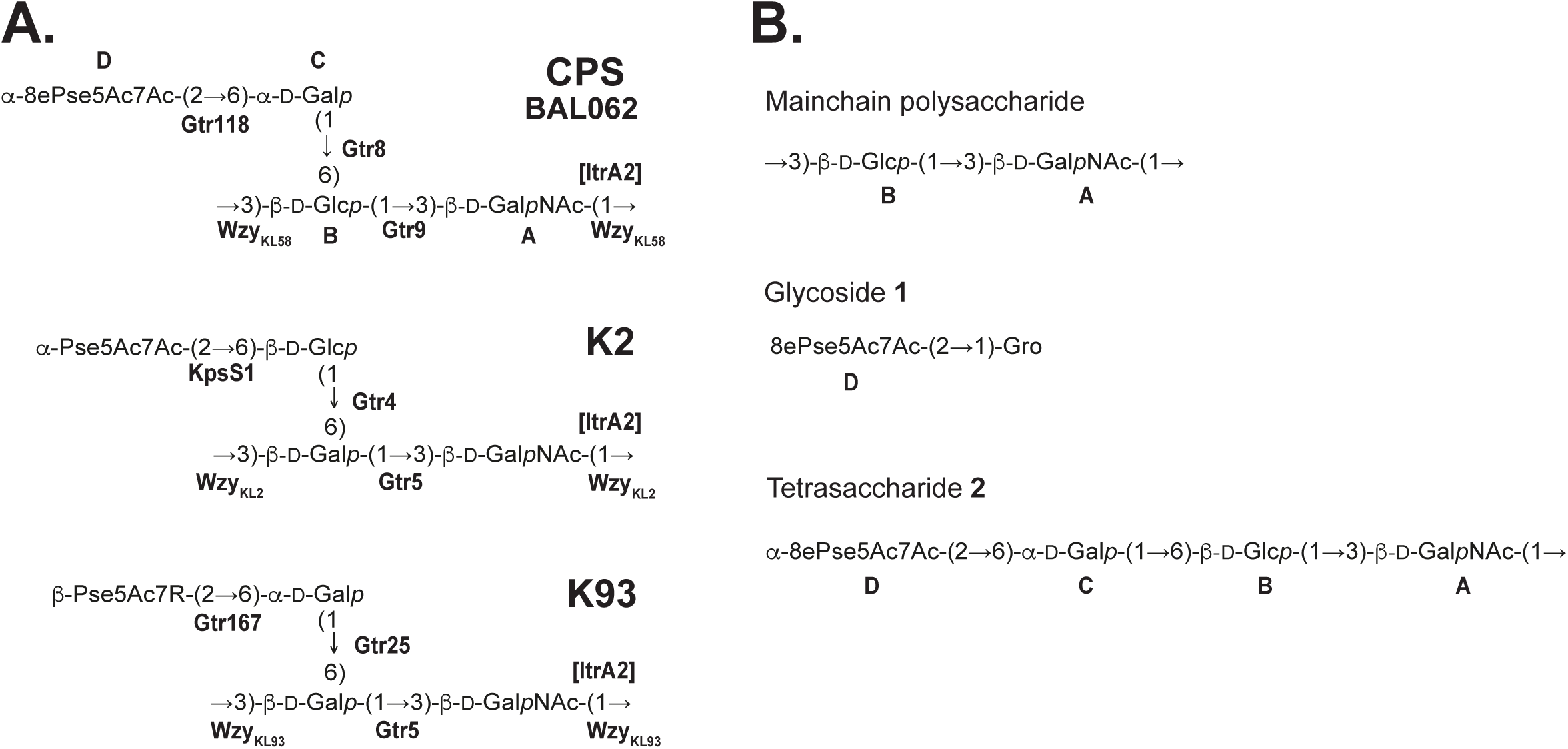
(**A)** Structure of the CPS produced by *A. baumannii* BAL062 compared with K2 (32, 33) and K93 (34). **(B)** Products derived by chemical cleavages of the BAL062 CPS. 8ePse5Ac7Ac indicates 5,7-diacetamido-3,5,7,9-tetradeoxy-d-*glycero*-l-*manno-*non-2-ulosonic acid (di-*N*-acetyl-8-epipseudaminic acid); Gro indicates glycerol.

**Table 2.** ^1^H and ^13^C NMR chemical shifts (δ, ppm) of the capsular polysaccharide produced by *A. baumannii* BAL062.

| Sugar | C3 or C1 | C4 or C2 | C5 or C3 | C6 or C4 | C7 or C5 | C8 or C6 | C9 |
| --- | --- | --- | --- | --- | --- | --- | --- |
|  | *H3ax, H3eq* | *H4 or H2* | *H5 or H3* | *H6 or H4* | *H7 or H5* | *H8 or H6* | *H9* |
|  | *or H1* |  |  |  |  |  |  |
| β-8ePseAc<sub>2</sub> **D** | 37.5 | 67.8 | 49.8 | 73.3 | 55.0 | 67.5 | 19.2 |
|  | *1.62, 2.50* | *3.90* | *4.18* | *3.87* | *3.89* | *4.37* | *1.08* |
| -6)-α-Gal **C** | 100.0 | 70.2 | 71.2 | 71.3 | 71.2 | 65.7 | |
|  | *4.96* | *3.83* | *3.97* | *3.83* | *3.99* | *3.57, 3.97* |  |
| -3,6)-β-Glc **B** | 106.0 | 74.7 | 85.8 | 70.0 | 75.6 | 67.8 | |
|  | *4.78* | *4.04* | *3.91* | *4.10* | *3.68* | *3.74, 3.80* |  |
| -3)-β-GalNAc **A** | 103.4 | 53.5 | 81.5 | 70.0 | 76.7 | 62.9 | |
|  | *4.51* | *3.40* | *3.66* | *3.59* | *3.62* | *3.74, 3.95* |  |

$^1\text{H}$ NMR chemical shifts are italicized.

The chemical shift for C6 of the higher sugar in the CPS (δ 73.3 ppm) is similar to the C6 chemical shift (73.0 ppm) of β-8ePse5Ac7Ac having the axial carboxyl group, but significantly different from that (70.3 ppm) of α-8ePse5Ac7Ac with the equatorial carboxyl group (36).

Therefore, 8-epipseudaminic acid in the CPS has the axial carboxyl group and is thus β-linked. The CPS from BAL062 therefore includes tetrasaccharide K-units with an 8ePse5Ac7Ac-(2→6)-Gal disaccharide branching from a disaccharide main chain composed of d-Glc*p* and d-Gal*p*NAc (Fig. 4A). The attachment of the side chain to position 6 of one of the main-chain components was confirmed by a glycosylation effect, that is a low-field position at 67.8 of the C6 signal of the d-Glc*p* monosaccharide that carries the side chain in the NMR spectra of the CPS, as compared with its position at ∼62-63 ppm in the spectra of the corresponding non-substituted monosaccharides.

### Assignment of encoded glycosyltransferases to linkages

The composition and topology of the BAL062 CPS is closely related to K2 and K93 types as predicted (Fig. 4A; and see above). As KL58 includes a gene encoding an ItrA2 transferase for initiating CPS synthesis by transferring d-Gal*p*NAc-1P to the lipid carrier (33), and a d-Gal*p*NAc residue is present in the CPS main chain, d-Gal*p*NAc was assigned as the first sugar (Fig. 4A). Hence, the β-d-Gal*p*NAc-(1→3)-β-d-Glc*p* linkage represents the bond between K-units that is likely formed by the Wzy_KL58_ polymerase (GenPept accession number SBS23904.1) encoded by KL58. Consistent with this conclusion, Wzy_KL58_ shares 84% amino acid (aa) sequence identity with Wzy_KL93_ (34) and 79% aa identity with Wzy_KL2_ (33), both of which form a similar β-d-Gal*p*NAc-(1→3)-β-d-Gal*p* linkage in the respective CPS (Fig. 4A). A further search of the BAL062 whole genome sequence did not detect any other Wzy gene candidates, hence Wzy_KL58_ encoded by the K locus was assigned to the β-d-Gal*p*NAc-(1→3)-β-d-Glc*p* linkage between units in the CPS structure.

The three glycosidic linkages in the K-unit are formed by glycosyltransferases encoded by the *gtr118*, *gtr8* and *gtr9* genes present in KL58 (Fig. 2). Gtr8 and Gtr9 have previously been found to form the respective linkages in an α-d-Gal*p*-(1→6)-β-d-Glc-(1→3)-β-d-Gal*p*NAc disaccharide in the K3-type CPS (37, 38). As the same segment is found in the BAL062 structure, Gtr8 and Gtr9 were assigned to these linkages (Fig. 4A). Hence, Gtr118 would be responsible for the β-8ePse5Ac7Ac-(2→6)-d-Gal*p* linkage in the side chain, and this is supported by Gtr118 sharing 82% aa identity with Gtr167 that forms a similar β-Pse5Ac7RHb-(2→6)-d-Gal*p* linkage in the K93 CPS (34).

### Distribution of the KL58 locus in A. baumannii genomes

In addition to the GC2 (n=23) and CC10 (n=6) KL58 isolates from the HTD outbreak, a search of 22,218 *A. baumannii* genomes available in the NCBI GenBank and non-redundant databases (as of 7^th^ February 2024) identified KL58 in a further 29 isolates (Fig. 5). These included ones from both clinical and environmental sources recovered over a period of two decades (2003 to 2023) from countries including the USA, Canada, China, Singapore, Germany, Poland, and Belgium. Despite the wide distribution, no further isolates from Vietnam or GC2 were detected. However, three isolates from either China (ST10=1; ST574=1) or Belgium (ST574=1) were CC10. The remaining genomes belonged to one of ten other STs or were non-typeable, and included either none or 1-2 resistance determinants (Fig. 5).

**Figure 5.**
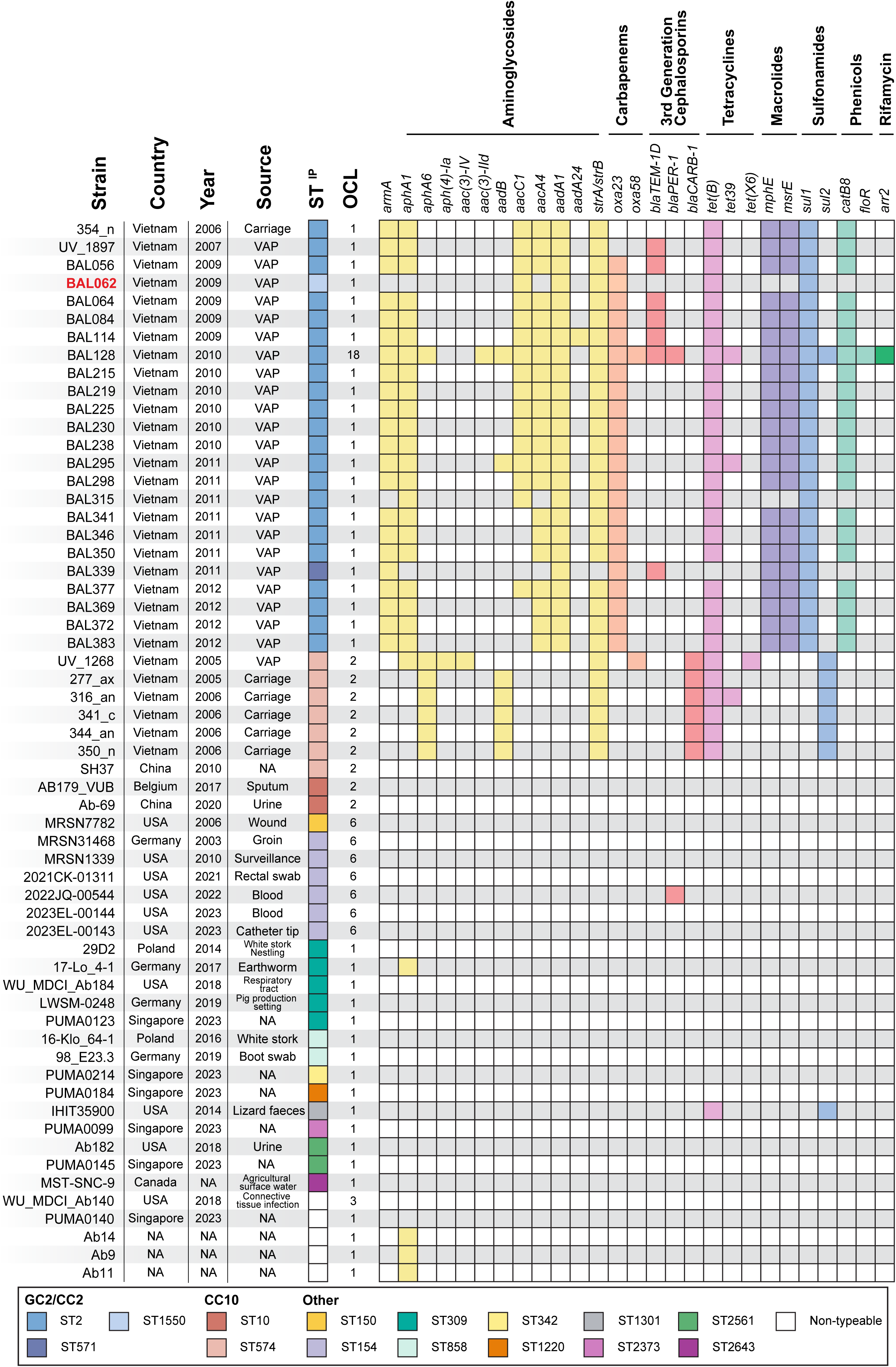
Distribution of KL58 in *A. baumannii* genome sequences. Colour scheme denoting STs in the Institut Pasteur scheme is shown below. NCBI accession numbers for isolates carrying KL58 are listed in Table S1.

## DISCUSSION

The use of contemporary nosocomial isolates such as BAL062, over ATCC reference strains isolated >70 years ago, has been recognized by many as key to obtaining data that is relevant to currently circulating clinical isolates (3–5). Although *A. baumannii* BAL062 is an important clinical GC2 reference isolate as it has been used in multiple experimental studies, some of its basic properties had not been reported. In this study, we report several key properties of the currently available form of this isolate and show that it has lost some of the resistance genes that would account for the phenotype of the original isolate (9).

As genetic manipulation relies on techniques that involve resistance markers, suitable strains would ideally be susceptible to one or more appropriate resistance markers. Hence, the previously unnoticed susceptibility to amikacin and kanamycin we have identified will be useful for future studies to replace difficult-to-use resistance markers for selection, including those for tellurite and hygromycin resistance, that are often used. In GC2 isolates, amikacin resistance can be directed by *armA* or *aacA4* located in AbGRI3, and these genes were found in the related lineage B GC2:KL58:OCL1 isolates from the HTD outbreak. However, as only a single IS*26* was found in the BAL062 chromosome at this location, it is likely that an IS*26*-mediated deletion (39) has occurred since BAL062 was first isolated. Likewise, the *bla_TEM-1D_*gene and *aphA1* kanamycin resistance gene was likely lost via an IS*26*-mediated deletion internal to AbGRI2 and such events have been reported previously (20). The loss of resistance markers that were likely present in the original isolate also highlights the importance of continually tracking the properties and potentially the genome sequences of isolates that are being used for experimental studies to ensure that they have not evolved in unexpected ways.

We further showed that BAL062 is a member of a discrete lineage of GC2 isolates, referred to as lineage B, from the HTD outbreak (15). A characteristic of this lineage is the presence of a Tn*2008VAR* transposon carrying *oxa23* that interrupts an acyl-CoA dehydrogenase gene in the chromosome. As a complete genome sequence was available for BAL062, we could accurately determine the precise location of the insertion via the identification of a 9 bp target site duplication (TSD) on either side of the transposon (Fig. 1A). This sequence was found to be different to that predicted previously, highlighting the importance of having a complete genome sequence available for clinical reference isolates.

Another characteristic of lineage B isolates and BAL062 is the presence of a KL58 sequence at the CPS biosynthesis K locus, which was found to be widely distributed and present in both clinical and environmental isolates. Granted the importance of the CPS and the influence of its specific structure on both virulence (40) and the application of alternate therapies such as monoclonal antibodies (41) and bacteriophage (42, 43), we used BAL062 to determine the K58-type structure. The non-2-ulosonic acid component of the K-unit was found to be 8ePse5Ac7Ac, the 8-epimer of Pse5Ac7Ac. This sugar was only recently discovered in the *A. baumannii* K135-type CPS (35), and it was later proposed that the genes responsible for conversion of Pse5Ac7Ac to 8ePse5Ac7Ac are located outside the K locus (44). Further work will be needed to identify the genetic determinant(s) for 8ePse5Ac7Ac for the K58 and K135 CPS forms. Nonetheless, the composition and overall topology of the CPS produced by BAL062 was found to be related to the K2 and K93 CPS as expected.

The structure correlated with the genetic annotation of KL58 gene cluster using agreed nomenclature is critical to future understanding of its role in or contribution to different phenotypes. In fact, a previous study that used the BAL062 TraDis library to identify genes involved in susceptibility to or tolerance of colistin (8) showed that that genes at the K locus as well as genes at the OC locus play a role. However, these genes were not identified as being in these locations.

Hence, the role of genes involved in synthesis of the outer core of LOS was not noticed. The potential role of CPS in colistin resistance or the involvement of K locus genes in synthesis of the LOS was also neither noticed nor explained, and further work will be needed to explain the role of the genes in colistin resistance. However, the location of genes in the K and OC loci was correctly identified in later studies (10, 11).

## MATERIALS AND METHODS

### Bacterial strain and antibiotic resistance profiling

*A. baumannii* isolate BAL062 was recovered in 2009 from a patient with ventilator associated pneumonia who was admitted to the ICU of the HTD in Ho Chi Minh City, Vietnam (6). The antibiotic resistance profile of BAL062 was determined as described previously (45).

### Bioinformatics analysis

The complete genome of BAL062 was downloaded from NCBI assembly accession number GCA_900088705.1 (chromosome: LT594095.1; plasmid: LT594096.1). KL and OCL sequences were identified by command-line *Kaptive v 2.0.7* using the current *A. baumannii* KL (28) and OCL (30) reference sequence databases. BLASTn was used to search 22,218 *A. baumannii* genomes in the NCBI GenBank and non-redundant databases (available as of 7^th^ February, 2024) for further instances of the KL58 sequence, and the associated metadata (country, collection year and source of isolation) were extracted from corresponding NCBI records and are compiled in Supplementary Table S1. For isolates reported in Schultz et al., draft genome sequences were assembled from short read data (SRA accessions listed in Table S1) using SPAdes (46).

Multilocus sequence typing (MLST) was performed using the *A. baumannii* Institut Pasteur scheme available at (https://pubmlst.org/bigsdb?db=pubmlst_abaumannii_seqdef). ResFinder *v 4.4.2* (47) was used to detect antibiotic resistance genes. The core-SNP maximum likelihood phylogeny was constructed using the Bactmap pipeline (https://github.com/nf-core/bactmap) with recombination removed using Gubbins (48), and the tree was visualised using iTOL (https://itol.embl.de/). Figures were created using EasyFig *v 2.2.2* (49) and annotated in Adobe Illustrator.

### Isolation of capsular polysaccharide

BAL062 was cultivated in 2×TY media overnight. Bacterial cells were harvested by centrifugation (10,000×*g*, 15 min), washed with and suspended in phosphate buffered saline. The suspension was cooled down to 4 °C, 0.2 volume of CCl_3_CO_2_H was added, cells were precipitated by centrifugation (15,000×*g*, 20 min), and two volumes of acetone were added to the supernatant. After intense shaking, a crude CPS preparation was separated by centrifugation (8,000×*g*, 20 min), dissolved in water, the pH value was adjusted to pH 8 by adding 1 M NaOH, the CPS was precipitated with acetone and separated by centrifugation as above, dissolved in distilled water and applied to a column (53 × 3.5 cm) of Sephadex G-50 Superfine (Healthcare). Elution was performed with 0.1% HOAc and monitored using a UV-detector (Uvicord, Sweden) at 206 nm to give purified CPS samples.

### Monosaccharide analysis

CPS samples (1 mg) were hydrolyzed with 2 M CF_3_CO_2_H (120 °C, 2 h). Monosaccharides were converted conventionally into the alditol acetates analyzed by GLC on a Maestro (Agilent 7820) chromatograph (Interlab, Russia) equipped with an HP-5 column (0.32 mm × 30 m) using a temperature program of 160 °C (1 min) to 290 °C at 7 °C min^-1^.

### Smith degradation

A CPS sample (54 mg) from *A. baumannii* BAL062 was oxidized with aqueous 0.05 m NaIO_4_ (1 mL) at 20 °C for 48 h in the dark, reduced with an excess of NaBH_4_ at 20 °C for 16 h. The excess of NaBH_4_ was destroyed with concentrated AcOH, the solution was evaporated, and the residue was evaporated with methanol (3 × 1 mL), dissolved in water (in 0.5 mL) and applied to a column (35 × 2 cm) of TSK HW-40. The degraded polysaccharide was eluted with aqueous 0.1% AcOH and hydrolyzed with 2 % HOAc (100 °C, 2 h) to give the β-8ePseAc_2_-(2→1)-Gro glycoside (5.2 mg) and a linear GlcNAc polymer (main-chain polysaccharide, 12 mg), which were isolated by gel-permeation chromatography on a column (108 × 1.2 cm) of TSK HW-40 in 1% HOAc.

### NMR spectroscopy

Samples were deuterium-exchanged by freeze-drying from 99.9 % D_2_O and then examined as solutions in 99.95 % D_2_O. NMR spectra were recorded on a Bruker Avance II 600 MHz spectrometer (Germany) at 60 °C. Sodium 3-trimethylsilylpropanoate-2,2,3,3-d_4_ (δ_H_ 0, δ_C_ −1.6) was used as internal reference for calibration. 2D NMR spectra were obtained using standard Bruker software, and Bruker TopSpin 2.1 program was used to acquire and process the NMR data. 60-ms MLEV-17 spin-lock time and 150-ms mixing time were used in ^1^H,^1^H TOCSY and ROESY experiments, respectively. A 60-ms delay was used for evolution of long-range couplings to optimize ^1^H,^13^C HMBC experiments for the coupling constant of *J*_H,C_ 8 Hz. ^1^H and ^13^C chemical shifts were assigned using two-dimensional ^1^H,^1^H COSY, ^1^H,^1^H TOCSY, and ^1^H,^13^C HSQC experiments (Table 2).

## Acknowledgements

We thank A/Prof Amy Cain (Macquarie University, Australia) for providing *A. baumannii* isolate BAL062, and Dr Stephanie Ambrose (University of Sydney, Australia) for technical assistance. NMR spectra were recorded in the Department of Structural Studies of N.D. Zelinsky Institute of Organic Chemistry, Moscow.

